# Evidence for auditory stimulus-specific adaptation but not deviance detection in larval zebrafish brains

**DOI:** 10.1101/2024.06.14.597058

**Authors:** Maya Wilde, Rebecca E. Poulsen, Wei Qin, Joshua Arnold, Itia A. Favre-Bulle, Jason B. Mattingley, Ethan K. Scott, Sarah J. Stednitz

## Abstract

Animals receive a constant stream of sensory input, and detecting changes in this sensory landscape is critical to their survival. One signature of change detection in humans is the auditory mismatch negativity (MMN), a neural response to unexpected stimuli that deviate from a predictable sequence. This process requires the auditory system to adapt to specific repeated stimuli while remaining sensitive to novel input (stimulus-specific adaptation). MMN was originally described in humans, and equivalent responses have been found in other mammals and birds, but it is not known to what extent this deviance detection circuitry is evolutionarily conserved. Here we present the first evidence for stimulus-specific adaptation in the brain of a teleost fish, using whole-brain calcium imaging of larval zebrafish at single-neuron resolution with selective plane illumination microscopy.

We found frequency-specific responses across the brain with variable response amplitudes for frequencies of the same volume, and created a loudness curve to model this effect. We presented an auditory ‘oddball’ stimulus in an otherwise predictable train of pure tone stimuli, and did not find a population of neurons with specific responses to deviant tones that were not otherwise explained by stimulus-specific adaptation. Further, we observed no deviance responses to an unexpected omission of a sound in a repetitive sequence of white noise bursts. These findings extend the known scope of auditory adaptation and deviance responses across the evolutionary tree, and lay groundwork for future studies to describe the circuitry underlying auditory adaptation at the level of individual neurons.

## INTRODUCTION

Animals receive a constant stream of sensory input, and detecting changes in this sensory landscape is critical to their survival. Our understanding of the neural circuits underlying this process across species is limited, especially at the level of individual neurons. In humans, one measure of brain activity following an unexpected change in sensory stimuli is known as mismatch negativity (MMN) ^1–3^. The mismatch refers to the elevated response to a deviant, unexpected stimulus (such as a different frequency tone), compared to a standard, predictable stimulus^1^. In the case of auditory stimuli, this signal is detected in the auditory cortex using electroencephalography or magnetoencephalography. Therefore, MMN in humans represents average activity in the auditory cortex, and the effect has not been recorded at the single-neuron level. A key feature of single-neuron activity underlying the MMN is stimulus-specific adaptation (SSA), where a neuron will decrease its response to a specific repeated stimulus, but this decrement does not generalise to all sounds^4^. While recognised as a crucial element of identifying deviant stimuli, it is debated whether SSA alone can represent true deviance detection, or whether a ‘pure’ mismatch effect requires SSA to be ruled out as a contributor ^4–6^. There is evidence for SSA in subcortical auditory structures in both humans ^7^ and rodents ^8,9^, notably the inferior colliculus and the medial geniculate nucleus of the thalamus ^4,10,11^. Subcortical SSA is also considered crucial for the MMN, but is typically weaker than cortical SSA^5^.

MMN is reduced in some neurological conditions such as schizophrenia and autism, and it therefore holds promise as a biomarker to aid understanding of neural circuitry underlying differences in sensory processing^12,13^. Developing MMN paradigms in animal models enables detailed investigation into the phenomenon and perturbations that affect it, leveraging their smaller brains to use techniques that are not feasible in human studies. An equivalent of the auditory MMN response is well established in rodents ^13–15^. MMN-like responses are also present in the equivalent of the auditory cortex in macaques^16^, cats^17^, pigeons^18^ and songbirds^19^, and there is preliminary evidence for auditory deviance responses in dogs^20^ and frogs^21^, though interpretation of these findings are confounded by lack of role reversal for the expected and unexpected sounds. MMN can also be elicited in other sensory modalities^1^, and indeed visual MMN has been characterized in mice using calcium imaging^22^

While the full complexity of the human MMN response may not be reflected in other species^14^, it stands to reason that the ability to detect changes in the sensory landscape would have arisen early in evolutionary time. It is not known to what degree this auditory deviance response is conserved across species, and it is not yet described in teleost fishes. Filling this gap in the evolutionary tree and identifying which homologous brain structures are required could help clarify whether these circuits evolved early in a common ancestor, or after the divergence of fish from mammals and birds.

Larval zebrafish are an appealing model for identifying whole-brain circuits underlying auditory deviance responses^23^. They are readily genetically modified, largely transparent, and have small brains, enabling the use of genetically-encoded calcium indicators for whole-brain imaging of activity at single-neuron resolution^24,25^. Additionally, they share many evolutionarily conserved and functionally homologous brain regions with other vertebrates, including mammals^26^.

At 6 days post-fertilization (dpf), zebrafish larvae show frequency-specific neuronal responses to sounds across a hearing range of at least 2 kHz^27^. While there is not currently evidence for predictive responses in the auditory domain, within the zebrafish visual system there is evidence for accumulation of sensory information and putative prediction error responses^28–31^. These studies use visual stimuli presented at a slower time course than is typically used for auditory MMN: many seconds, compared to auditory MMN stimuli which are typically tens of milliseconds long, with hundreds of milliseconds interstimulus interval. Nevertheless, these studies provide evidence that larval zebrafish exhibit a degree of sensory predictive activity. This predictive activity, coupled with their ability to distinguish different sounds, are necessary precursors to auditory deviance responses. We therefore designed an auditory stimulus paradigm to elicit deviance responses in larval zebrafish, and simultaneously imaged whole-brain neuronal activity at single-cell resolution.

## METHODS

### Animals

Adult zebrafish (*Danio rerio*) were maintained on a TLN background at a density of 10-15 fish per litre, at approximately 28.5°C with a light/dark cycle of 14/10^32^. We used *HuC:H2B-GCaMP6f* fish with targeted expression of the fluorescent calcium indicator GCaMP6f in the nuclei of neurons^33^.

Embryos were raised in petri dishes containing E3 media (distilled water with 10% Hanks solution, consisting of 137mM NaCl, 5.4mM KCl, 0.25mM Na2HPO4, 0.44mM KH2PO4, 1.3mM CaCl2, 1.0mM 654 MgSO4 and 4.2mM NaHCO3 at pH 7.2)^32^ in an incubator kept at 28.5°C with a 14/10 hour light/dark cycle. Zebrafish housing, breeding, larval maintenance, and experiments were performed with approval from the University of Queensland Animal Ethics Committee (IMB/237/16/BREED and SBS/341/19).

### Calcium imaging

Larval zebrafish aged 6 dpf were embedded in 2% low melting point agarose and mounted in custom 3D-printed 24×24 mm experimental chambers^27,34^. Chambers consist of corner posts with 20 mm square coverslips affixed using waterproof glue (Liquid Fusion Clear Urethane Adhesive) to form the chamber walls. A speaker affixed to the back wall delivered acoustic stimuli, such that the air-water interface was eliminated to improve stimulus delivery^27^. The chamber was then filled with E3 media. Fish were left to settle in the chamber for at least 60 minutes to prevent z-drift during acquisition.

Whole-brain fluorescence calcium imaging was conducted with a custom-built selective plane illumination microscope (SPIM), as described previously^35–38^. We used two light sheets, from the anterior and one side of the fish, to illuminate the full brain (Figure 1A). Sheets were scanned at 25 10µm steps throughout the dorsoventral axis of the animal at a rate of 4 volumes per second.

**Figure 1:**
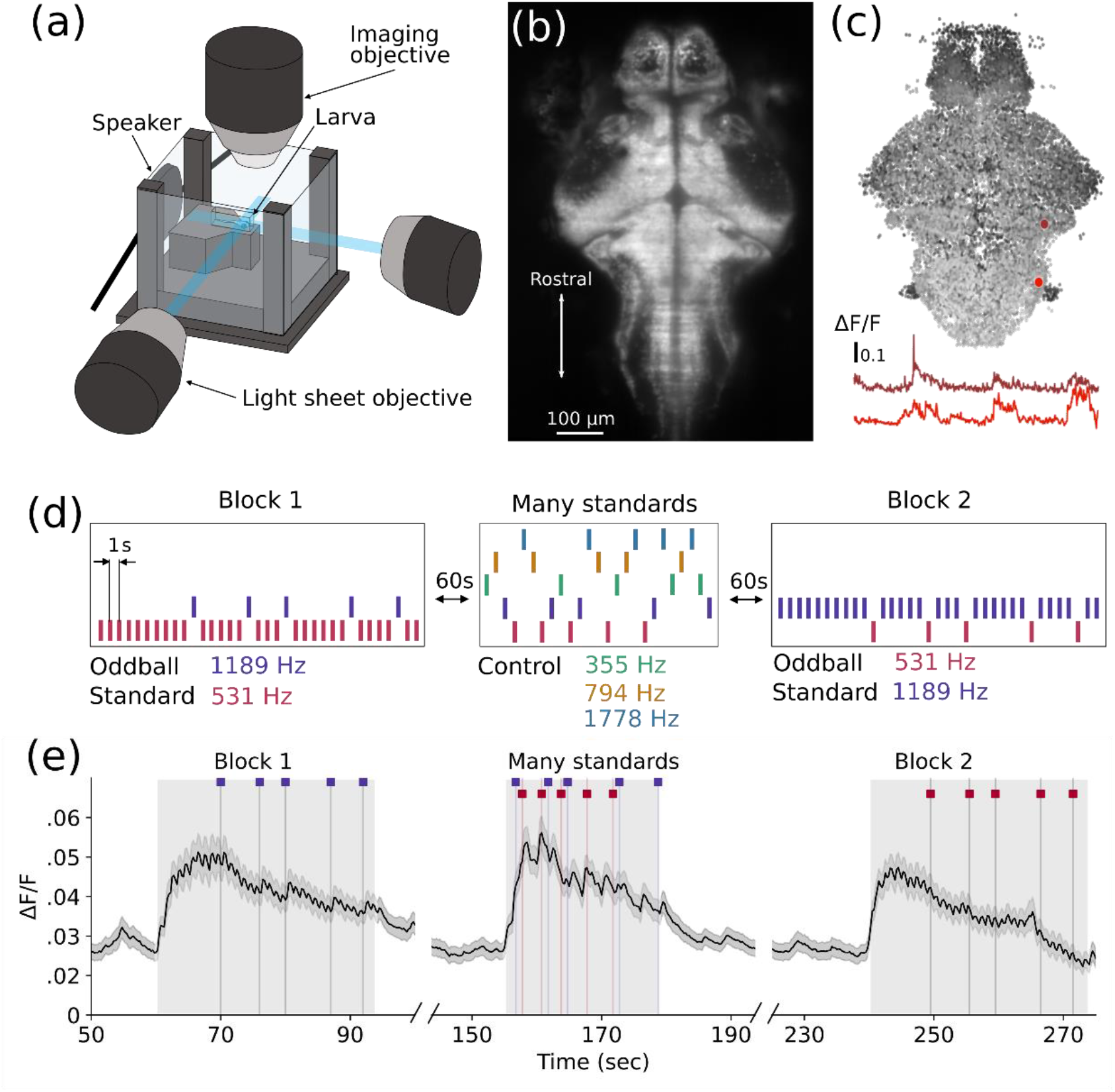
Calcium imaging of responses to auditory stimuli. **A)** Experimental set-up. A larva mounted in low melting point agarose is illuminated by two perpendicular sheets of light. Acoustic stimuli are delivered by a speaker affixed to the back wall. Calcium activity is captured with a water-immersion objective. **B)** Example single plane of a zebrafish larval brain expressing nuclear localized GCaMP6f under the control of the HuC promoter. Image was averaged over the entire recording after motion correction. **C)** Visualisation of all neurons segmented from a single larva, coloured by depth (black is most ventral, white most dorsal). Example neurons are highlighted in red, with their respective calcium activity traces over time represented below. **D)** Stimulus train. Experimental tones of 531 Hz (pink) and 1189 Hz (purple) each act as the standard and deviant stimulus in different blocks. Control tones of 355 Hz, 794 Hz and 1778 Hz enable each frequency to be presented at the same probability in the many standards control block. **E)** Average ΔF/F traces over all neurons (n = 186,675) from all fish (n = 12) for each block of the experiment. Grey boxes indicate the timings of the stimulus blocks. Stimulus times for the experimental tones when not standard are illustrated above each graph.

Both the image acquisition and delivery of acoustic stimuli were controlled using Micro Manager software (version 1.4)^39^. An example image of a single plane is shown in Figure 1B.

### Image processing pipeline

Individual regions of interest (correlating to neuronal nuclei) were segmented using Suite2p^40^, and their fluorescence traces over the course of the experiment were extracted (Figure 1C). The motion-corrected 3-dimensional mean image of each fish was then registered to the Zbrain reference larval zebrafish brain atlas^41^, using Advanced Normalization Tools (ANTs)^42^. These steps were completed within a custom data processing pipeline run on the high-performance computing cluster at the University of Melbourne. These data were then extracted and sorted in MATLAB (MathWorks, 2022b). Masks from the Zbrain atlas were used to remove any regions of interest detected in the eyes or otherwise outside the brain.

### Acoustic stimuli

Sounds were delivered to the fish with a mini speaker (Dayton Audio DAEX-9-4SM Skinny Mini Exciter Audio, Haptic Item Number 295-256), driven by an amplifier (Dayton Audio DA30 2 × 15W Class D Bridgeable Mini Amplifier), as previously described^27^.

#### Oddball tones

The 5 frequencies were chosen to be equally spaced on a logarithmic scale within the hearing range of larval zebrafish^27^. Sound stimuli were 100ms in length, with 2ms on and off ramps in volume. The volume was set to −6 dBFS (decibels digital Full Scale), equivalent to 96 dBSPL (decibels sound pressure level) when measured in a chamber not filled with E3^34^. The stimulus paradigm for these experiments followed a ‘flip, many standards, flop’ paradigm (Figure 1D)^15^. The experimental blocks consisted of 25 stimuli, of which 5 (20%) were deviant. Blocks 1 and 2 were preceded by 10 repetitions of the standard stimulus to establish the predictable sequence. Each block was separated by 60s, and stimuli within a block were presented with an interstimulus interval of 1 second. All larvae (n = 12) received the same pseudorandom stimulus train, with the same order of stimulus blocks, for a total of 95 stimuli.

#### Loudness curves

Nine frequencies were chosen to be equally spaced on a logarithmic scale within the hearing range of larval zebrafish, each played for 100ms at 4 volumes: 0, −6, −12, and −18dBFS.The order of these was randomised into two different sequences, and each fish (n = 18) received one of these.

#### Silent gaps

Acoustic stimuli were 100 ms white noise bursts at −6 dBFS, with 2ms on and off ramps, and the deviant event was the absence of this stimulus. These were delivered with an interstimulus interval of 500ms. Thirty white noise bursts were delivered to establish the sequence of sounds, then 400 stimulus timepoints, 40 (10%) of which were gaps (white noise was not delivered). At least 2 white noise bursts were delivered between each instance of a gap. All fish (n = 9) received the same pseudorandom stimulus sequence, totalling 390 sounds and 40 gaps.

### Analysis

Auditory regressors were generated from stimulus onset times convolved to a synthetic calcium signal that mimics the rise and decay dynamics of our fluorescent probe. Regressors for motor activity in each fish were similarly generated using the motion correction output from Suite2p. Cells were considered significant for a regressor if they met two criteria: 1.) a p value < .001 divided by the total number of cells in the experiment across animals and 2.) An R^2^ value greater than the 99th percentile of all cells, for that regressor.

Permutation clustering analysis was performed using k-means clustering of ΔF/F for all cells that passed the threshold for any of our previously described regressors, including fictive controls (8,306 cells total). Each iteration consisted of 30 clusters with randomly initialized centres, for a total of 33,000 unique clusters. The mean fluorescence of each cluster was correlated to both our oddball and offset control regressors. Clusters with an R^2^ value to the oddball regressor greater than the 99^th^ percentile (0.350, for total of 494 cells spanning 60 clusters) were then pooled to improve coverage across animals and used for subsequent calculations.

When applicable, paired t-tests were used for multiple comparisons with an alpha cutoff of 0.05 adjusted by a Šidák correction for multiple comparisons. Error is reported throughout as standard error of the mean.

Analyses were performed using custom software written in Julia 1.7.2 using GLM.jl (http://github.com/stednitzs/oddball/).

#### Oddball tones

The change in fluorescence over the baseline fluorescence (ΔF/F) was calculated using a sliding window of 201 timepoints (50.25 seconds) and a smoothing window of 7 timepoints.

#### Loudness curve

Auditory responsive neurons were identified using a regressor to the timing of each sound stimulus. The mean amplitude of the z-scored responses of these neurons to each frequency at each volume was calculated for each fish to produce the loudness curve. Z-scored responses were used to directly compare between neurons of different brightnesses to better capture the activity of dim cells that respond weakly to perceptually quiet stimuli. We used a linear mixed model to compare response amplitudes, with each fish as a random effect to control for repeated measures.

#### Silent gaps

To better correct for sustained, elevated calcium signals induced by closely spaced white noise stimuli, z-scored responses were used. Regressions were performed only on time periods during stimulus presentation to account for bleaching at the start of recording. We calculated the mean ΔF/F for gap responsive cells during the gap, and performed a linear regression using the number of preceding white noise stimuli as a predictor.

## RESULTS

### Auditory responses

The experimental design followed a flip-many standards-flop paradigm, consisting of three stimulus blocks (Figure 1D)^15^. This design enables us to compare responses to the same tone when it is the standard or deviant stimulus, and controls for different sensitivity to each frequency. The many standards block is an additional control, where three other frequencies are included such that there is no expectation set by a predictable sequence, but the likelihood of each stimulus is equal to the deviant tone in the predictable sequences.

Using a regression-based approach, we identified individual cells in animals that responded to specific aspects of the auditory stimuli. First, we classified cells as broadly auditory responsive (responding to all sounds) (Figure 2A-B). This approach permitted us to identify cell populations corresponding to previously identified brain regions in the zebrafish auditory processing pathway, including the octavolateralis nucleus, torus semicircularis, cerebellum, thalamus and telencephalon (Figure 2A, Figure S1A-B)^27,43^. These broadly auditory cells respond to reference, many standards, and oddball tones, regardless of the preceding stimulus presentation (Figure 2B). The spatial distribution of these neurons resembles previously identified distributions of pure tone-responsive neurons in the larval zebrafish brain^27^.

**Figure 2:**
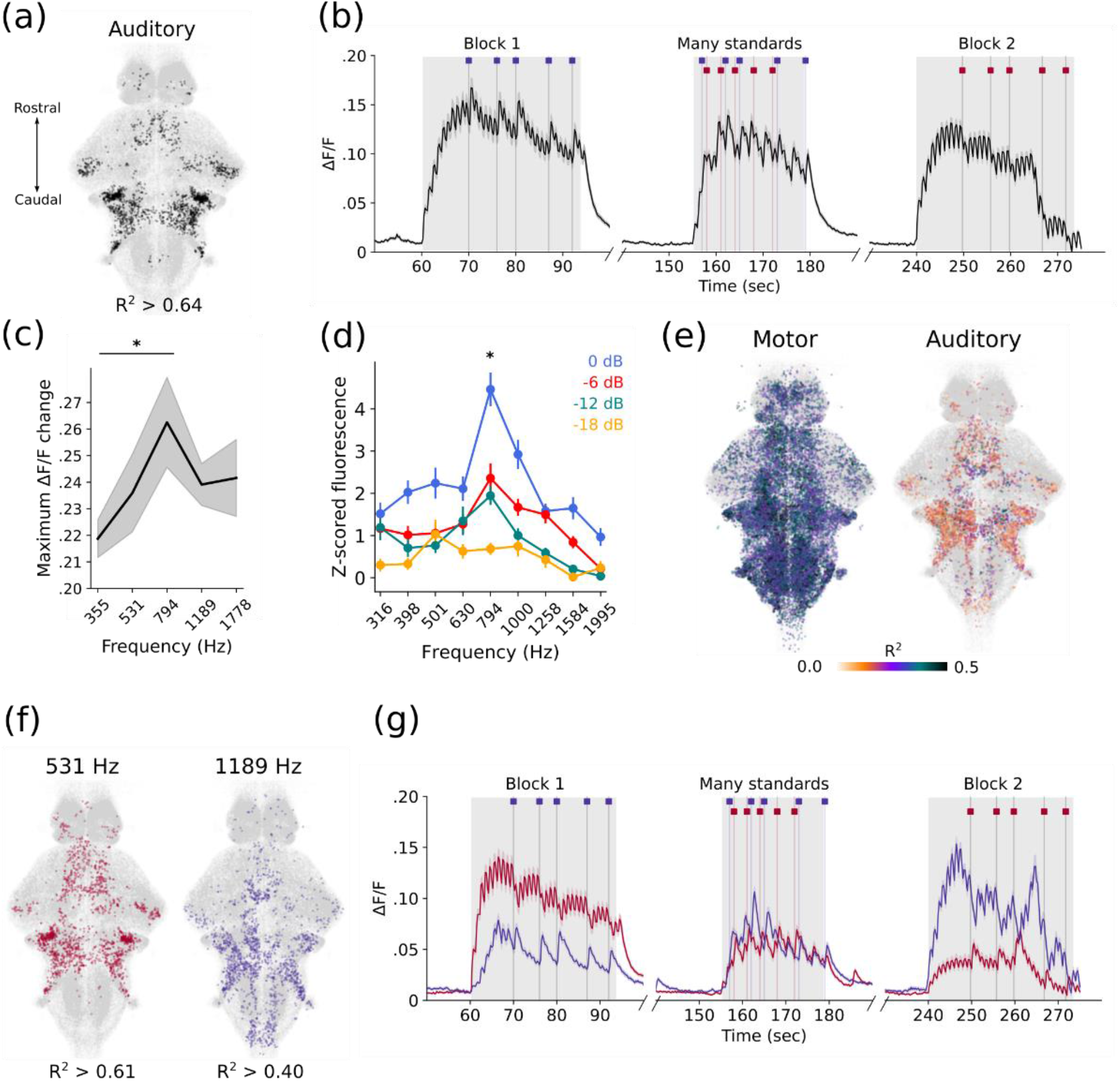
Auditory specific responses. **A)** Spatial distribution of generally auditory responsive neurons (black). R^2^ value is the threshold to attain 99^th^ percentile of auditory neurons **B)** Mean trace of all neurons passing the criteria for classification as generally auditory responsive. Timings of the 3 experimental blocks are indicated by grey shading behind the trace. **C)** Amplitude of the responses of auditory neurons to each frequency in the many standards control block. Grey shading represents standard error of mean. * = p <0.01 for 794 vs. 355 Hz. **D)** Equal loudness contours for larval zebrafish, separate dataset. Dots indicate means over 18 fish and bars indicate SEM. Note frequencies are evenly spaced within a logarithmic distribution. Significant effects of volume (p < 0.001) and frequency (p < 0.005), but no significant interaction (p = 0.236). * = p < 0.01 **E)** Correlation to motor activity. Left indicates distribution of motor responsive neurons, right indicates the motor correlation values of neurons that were classified as auditory responsive. Colour of dots represents the R^2^ value of the correlation to the motor activity. **F)** Spatial distribution of frequency specific neurons: those preferentially responding to 531 Hz (pink), and those preferentially responding to 1189 Hz (purple), from across all animals. Populations defined by responsiveness to all presentations of that frequency across blocks. R^2^ values are the thresholds to attain 99^th^ percentile of each of these populations. **G)** Mean traces in each block of the experiment of all 531 Hz-selective neurons (pink), and 1189 Hz-selective neurons (purple). Timings of the 3 experimental blocks are indicated by grey shading behind the trace. Timings of the deviant stimuli are indicated by the coloured dashes above the trace.

However, we noticed that the response amplitude in these auditory neurons was lower for 531 Hz tones than 1189 Hz tones in the first block, despite being presented at the same intensity (Figure 2B). This could be a result of either adaptation or salience differences between these frequencies. We computed the maximum change in ΔF/F for each frequency in the many standards block only, finding that on average auditory cells exhibit larger calcium responses to frequencies between 531-1189 Hz (Figure 2C). This led us to question if larval zebrafish exhibit equal-loudness contours similar to other animals, where frequencies in the middle of the hearing range are perceived as louder than those towards the edges, even when presented at equal intensity^44^. We therefore collected another dataset to establish the landscape of equal-loudness contours, using nine logarithmically-spaced frequencies across the hearing range of larval zebrafish^27^. Indeed, the curve produced implies increased loudness for frequencies toward the centre of the hearing range compared to the higher and lower end (Figure 2E). We analysed this loudness curve statistically, and found main effects for both volume and frequency (p < .001 and .005 respectively), and no overall interaction effect (p = .236.). At the volume used in the main experiment (−6 dBFS), responses to 794 Hz are significantly higher than frequencies on the periphery (p < 0.01). These curves are evidence that 1189 Hz tones have a greater effective loudness, and therefore salience, than 531 Hz tones across different volumes for larval zebrafish.

Although we recorded animals under a restrained preparation that did not permit the recording of tail movements, we inferred movement attempts from the motion correction derived from our processing pipeline (Figure S1C). Movement efforts are highly correlated to cells in the hindbrain, corresponding to motor command regions (Figure 2E)^45^. A proportion of auditory-responsive cells are also correlated with movement events, indicating movements often occur simultaneously to an auditory stimulus, but not consistently enough to suggest the requirement for motion correction arises from vibration of the chamber during stimulus presentation (Figure 2E). We find no consistent overall differences in the behavioural response rate from the two frequencies most relevant to our experimental paradigm (Figure S1D).

Given that some auditory cells may be frequency-selective, we further defined two populations of neurons that preferred each of the experimental frequencies (Figure 2F-G). These populations are similarly spatially distributed throughout the brain (Figure 2F), and present in all animals in our experiment (Figure S1E-F). The mean activity traces of each of these populations indicate they also respond to the non-preferred frequency, but to a lower degree than their chosen frequency (Figure 2G). In blocks where the non-preferred stimulus is the standard tone, there is reduced calcium accumulation, and lower overall activity, while the activity within the many standards control block is comparable. This effect is consistent with cells that respond preferentially to specific frequencies, as the number of stimulus presentations in the many standards block is equivalent for both tones.

### Stimulus specific adaptation

We observed that by the second block, responses to the 531 Hz deviant tone were suppressed relative to the first block (Figure 2B, 2G), which could be explained by stimulus-specific adaptation. This adaptation is regarded as a key component underlying auditory deviance detection^4^. We measured the mean change in ΔF/F at each stimulus presentation for broadly auditory cells, and compared this to the first 5 presentations of each stimulus in all three blocks (Figure 3A). We find that responses decay over repeated presentations, even when the tone is unexpected. We also included the 5 standard presentations that directly precede an oddball to account for the total number of preceding stimuli, and found that these responses were decreased after many repetitions, consistent with adaptation. In contrast, the responses to the deviant frequency in the first block remain elevated during the many standards, suggesting the 5 preceding presentations of that frequency were insufficient to induce adaptation. Conversely, in the last block, responses to that frequency as the standard are initially elevated but then decrease to the same level as the adapted frequency. If the activity of these neurons also encoded the unexpected nature of a stimulus, the responses are expected to be greater when it is the oddball rather than the predictable standard preceding an oddball. However, this is only the case in the first block, where the oddball was not previously adapted (Figure 3B).

**Figure 3:**
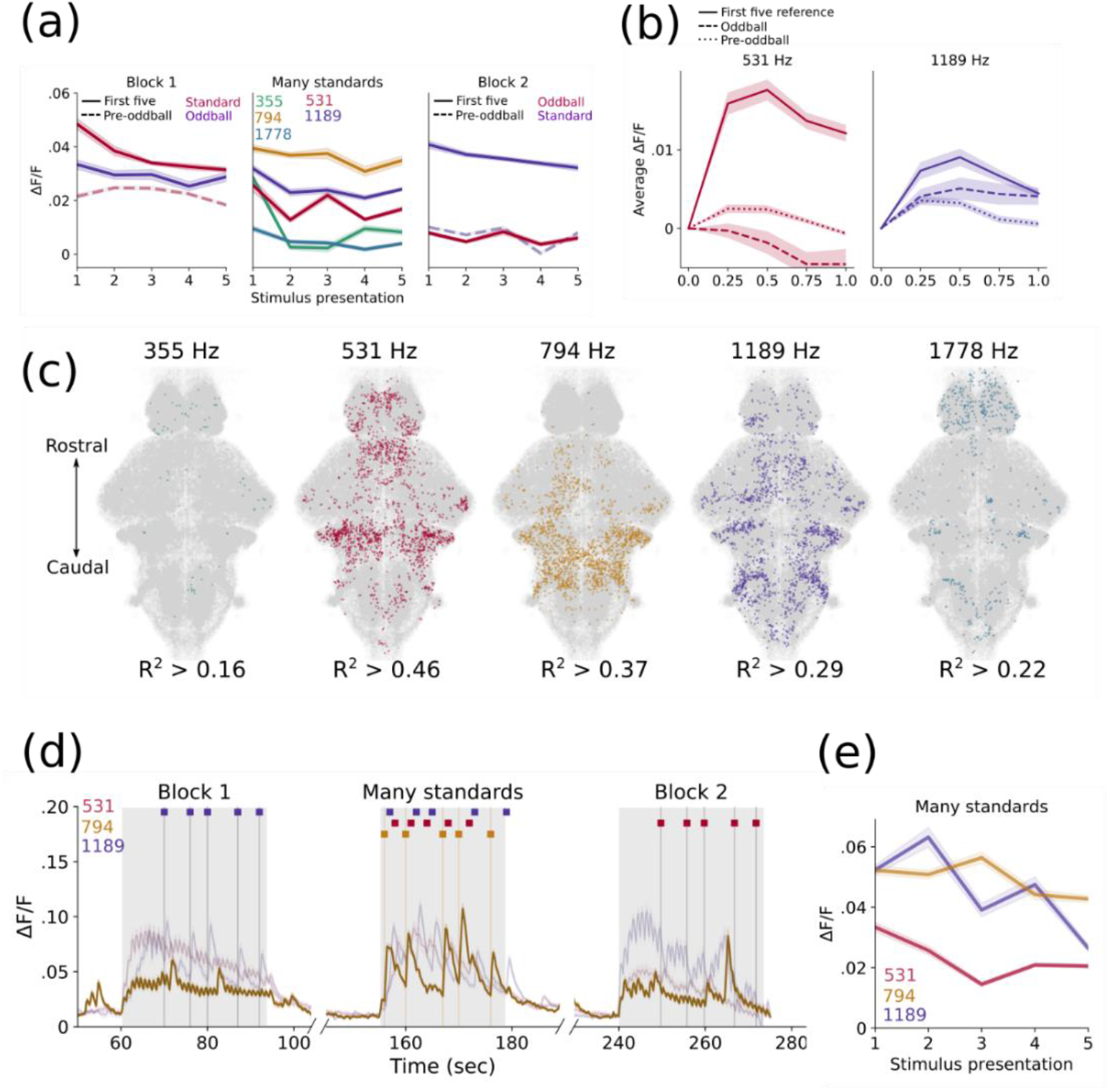
Stimulus specific adaptation. **A)** Mean response amplitudes of all auditory-classified neurons to specific stimulus presentations. In block 1 and 2, solid lines represent the first 5 presentations of each frequency (as either standard or oddball), and the dotted line represents the 5 presentations of the standard stimulus directly preceding an oddball. In the many standards block, the mean responses to 5 presentations of each frequency are shown. Shading represents standard error of mean. **B)** Mean traces of frequency selective neurons to the first 5 reference tones of each frequency, the 5 oddball presentations of each frequency, and the 5 instances where each frequency directly precedes an oddball when it is the standard. Shading represents standard error of mean. **C)** Spatial distributions of 5 populations of frequency-specific neurons defined by regressions to their presentation during the many standards control block. R^2^ values represent the threshold to attain the 99^th^ percentile of responses. **D)** Mean traces throughout the whole experiment of the 794 Hz-preferring population (dark yellow). Fainter lines for the 531 and 1189-preferring populations are included for reference. Timings of the 3 experimental blocks are indicated by grey shading behind the trace. Timings of the two experimental frequencies when infrequent and the preferred 794 Hz tone are indicated by the coloured dashes above the trace. **E)** Average responses of each of the 3 middle frequency preferring populations to each of the 5 presentations of their preferred frequency during the many standards block. Shading represents standard error of the mean.

We then used the many standards control block alone to define populations of frequency-selective cells to assess whether off-target frequencies produced adaptation in each population and partially control for order effects (Figure. 3C). We did not detect populations that correlate well with tones at the extreme ends of the hearing range, consistent with the weak responses detected in our analysis of auditory cells (Figure 2C) and previous studies^27^. We therefore focused on the three middle frequencies, and found that each population also responds to the other frequencies, despite preferring its assigned frequency (Figure 3D). However, the population that prefers the initial standard frequency (531 Hz) are more adapted by the many standard blocks than other populations (Figure 3E). Interestingly, the responses of the 794 Hz neurons are similar in block 1 and block 2, suggesting the number of preceding off-target stimuli doesn’t affect future responses to off-target stimuli, therefore adaptation is specific to the preferred frequency (Figure 3D). As expected, 1189 Hz neurons maintain high response amplitudes until late in block 2, despite many preceding off-target frequency stimuli (Figure 3D).

If these activity patterns are explained by SSA, we would expect the rate and extent of adaptation to decrease with longer interstimulus intervals^5,38^. To test this, we used an identical protocol with 2 and 3 second interstimulus intervals and found longer interstimulus intervals produce less pronounced SSA (Figure S2A-D).

### Oddball tone

Given the response amplitude of frequency-selective neurons was not reliably higher to oddball stimuli than standard stimuli (Figure 3B), we instead looked for a population of neurons that responds specifically to unexpected sounds. Using regressors to the timing of deviant sounds in blocks 1 and 2, we then identified cells that selectively respond to oddball stimuli in both blocks (Figure 4A-C). As a control we included two additional variables, consisting of the same number of fictive stimuli locked to the presentation of actual auditory pulses: 1.) random, where fictive stimuli can occur at any time during the stimulus train, including during oddball presentations, and 2.) offset, where fictive stimuli are constrained to occur only when oddball stimuli are not presented (Figure 4A). These controls allowed us to rule out cells that would be randomly correlated simply by the periodic nature of their firing rate.

**Figure 4:**
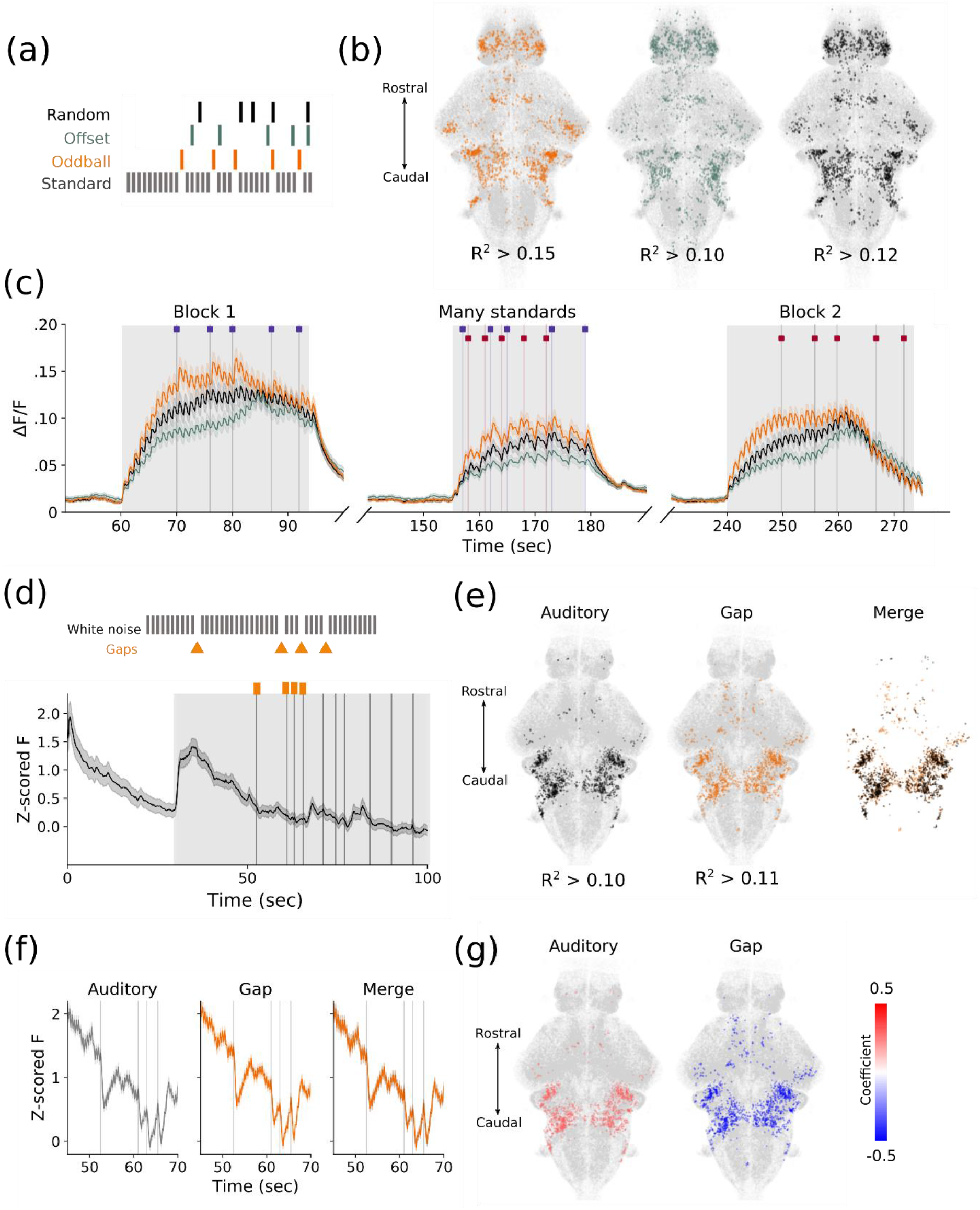
No evidence for oddball-specific responses. **A)** Example of regressor timings to the oddball (orange), and the offset (cannot be aligned with oddball, green) and random (can occur at any time, black) controls. **B)** Distribution of oddball responsive (orange), offset control (green) and random control (black) neurons. R^2^ values are the threshold cutoff for the 99^th^ percentile of responses. **C)** Mean traces of neurons classified as high fidelity to the oddball, offset, and random stimulus timings. D-G: Silent gap experiments. **D)** Example of early part of the stimulus train. The standard stimulus is a white noise burst, and gaps in this sequence are the deviant stimuli (orange). Below is the Z-scored fluorescence for all cells in the early part of the experiment, average across all fish (n = 9). Vertical lines indicate gaps in the stimulus train. Shaded region indicates the portion of the stimulus train illustrated above. **E)** Neurons that regress significantly to the white noise bursts, vs the gaps, and their overlap. R^2^ values are the threshold cutoff for the 99^th^ percentile of responses. **F)** Average Z-scored fluorescence for cells that regress significantly to the white noise burst, the gap, and their overlap. The timepoints for these examples match those of the inset in (D). **G)** Regression coefficients for auditory and gap responsive cells, indicating positive and negative coefficients respectively.

The cells in each of these three populations had similar spatial distributions (Figure 4B) and were all broadly auditory responsive (Figure 4C). While we could identify a population of neurons that correlated to the specific timing of the oddball tones, we could similarly find populations with an equivalent correlation to random subsets of stimulus timings. A higher proportion of cells in the telencephalon responding preferentially to oddball tones (33.4% ± 1.4) than in the offset (4.38% ± 0.9, p < 0.002) and random conditions (22.2% ± 1.3, p < 0.001). This effect may be an artifact of high rates of spontaneous activity in the telencephalon, and cannot be conclusively identified as a deviance-selective population using a regression approach.

We considered the strictness of our regression criteria could hinder our ability to detect deviance responsive cells. Therefore, we performed a permutation clustering analysis with 100 iterations of k-means using randomly initialized centres, which should capture even rare patterns of activity. We subsequently correlated the cluster average to our oddball regressor to identify any clusters characterized by deviance responses. While we were able to identify clusters that were more highly correlated to unexpected stimuli than others (60 clusters with R^2^ > 99^th^ percentile compared to a mean of 0.158 across all clusters), these consisted of either few cells or were poorly distributed across animals (Figure S2E). We pooled all clusters with R^2^ > 99^th^ percentile and computed the average response to oddball and reference tones in both blocks, finding no significant differences between them, in contrast to the expected behaviour of deviance-detecting cells (Figure S2F). We further note similar correlations to the fictive offset control, and therefore conclude that these correlations are spurious.

### Silent gap

We questioned whether the lack of clear deviance-specific responses was due to the masking of salience and order effects. We therefore used a paradigm where the deviant stimulus was the absence of a predictable sound, so that any response at that time could represent predictive activity (Figure 4D). We identified neuronal populations that responded specifically to the white noise sound stimuli or the gap and found nearly completely overlapping spatial distributions and activity (Figure 4E-F). Indeed, responses to gaps occurred only in cells that were also auditory sensitive, and were negatively correlated with gaps (Figure 4G), indicating a drop in calcium consistent with auditory cells. The magnitude of this drop was not correlated to the number of preceding sounds, which would have indicated it was related to how unexpected the gap was (Figure S2G). Finally, we did not find a population of neurons whose activity correlated positively with the gaps, suggesting that zebrafish larvae do not exhibit MMN-like responses to the unexpected absence of auditory stimuli.

## DISCUSSION

### Stimulus specific adaptation in larval zebrafish

We found that the two frequencies of sound used in the experiment had varying salience for the fish (Figure 2C-D), and each recruited frequency-specific populations of neurons (Figure 2F-G).

We also demonstrated here an effective loudness curve for larval zebrafish (Figure 2D), which will aid the design of future research into auditory responses in larval zebrafish. These results confirmed that the larval nervous system has the capacity to distinguish between distinct frequencies. We observed a decrease in the response amplitude across the course of the experiment, representing adaptation (Figure 2B). This decrease is unlikely to be due to photobleaching, as the responses recover after the brief break between blocks of stimuli.

Within the general population of auditory neurons, we found that the degree of adaptation to one frequency did not generalise to the other frequency (Figure 3A-B). While many stimulus repetitions cause decreased responses to a given sound, the same reduction is not observed when the sound is of a different frequency (Figure 3A).

To understand if this is a property of neurons that respond to any tone or can also be observed in frequency selective cells, we measured stimulus-specific adaptation within populations of neurons that prefer a specific frequency (Figure 3C-E). As expected, we did not find neurons selective for frequencies at the far ends of the hearing range, consistent with the profiles of frequency-specific populations in a previous study^27^. These neurons were not entirely frequency selective, rather they exhibit smaller responses to off-target frequencies (Figure 3D). However, these weak responses to other frequencies do not impact the subsequent response to the preferred frequency. For example, the 1189 Hz cells do not decline in activity until the final block, when it is consecutively repeated. In contrast, the 531 Hz population is already adapted during the first block, when it is the standard tone, and their responses remain low throughout the subsequent blocks.

These results suggest stimulus-specific adaptation in the auditory domain in larval zebrafish, a critical requirement for the ability to detect auditory deviance responses^4–6^. The neurons showing SSA are located throughout the fish auditory system: in the octavolateralis nucleus (homologous to the mammalian cochlear nucleus), the torus semicircularis (homologous to the mammalian inferior colliculus), the cerebellum, thalamus and telencephalon. These auditory regions represent the zebrafish homologues of the subcortical auditory pathway in mammals^43^, which exhibit SSA as a building block underlying auditory deviance detection in higher order brain structures^4,5^. Our findings are therefore consistent with the idea that SSA evolved in a common ancestor, and that its fundamental mechanisms may be conserved between zebrafish and other species.

SSA, as described here, is a crucial element underlying the ability to detect auditory deviance^4^. Further, a deviance-specific response can be established by the inclusion of both predictable blocks and the unpredictable many-standards control, where the response to a given frequency is elevated only when it is unexpected^14^. Studies in rodents and pigeons successfully identified responses meeting these criteria^14,15,18^. However, efforts to identify an equivalent of MMN in songbirds and frogs failed to find similarly deviant-specific responses that cannot be explained by stimulus salience or SSA^19,21^. From our findings, the larval zebrafish fit into the latter category. This does not exclude the possibility that deviance-specific responses exist in these animals, but perhaps the optimal experimental paradigm has not yet been tested. In our case, larval zebrafish do not have fully developed nervous systems and true deviance detection may arise later in development.

### Future directions for identifying deviance responses

A challenge in designing an auditory deviance paradigm for larval zebrafish is the lack of an expected behavioural output. Indeed, although we detected some behavioural responses to auditory stimuli despite larvae being physically restrained, these did not correspond specifically to deviant stimuli. While it is established that zebrafish possess distinct cell populations that are frequency-selective^27^, and we recapitulate those findings here, the ethological relevance of this is unclear. For example, some species of teleost fish employ auditory communication, but this is not reported in zebrafish^46,47^. Another possibility is that frequency selectivity is a nascent property of the larval brain that will become ethologically relevant as the animal develops further. Identifying the functional purpose of frequency selectivity could inform an experimental paradigm that induces more detectable and consistent deviance responses.

One aspect of deviance detection in mammals is a response to the lack of an expected stimulus. Such an experimental design also eliminates the salience and order effect confounds seen in the experiment with pure tone stimuli, but responses to missing stimuli are more difficult to characterise^48^. With this approach, any positive deflections in the activity at the time of the gap between predictable sounds are more likely to be a prediction error, rather than a preference for a given frequency. However, the only neurons we found with significant correlation to our regressor to the timings of the gaps were negatively correlated, consistent with the expected behaviour of auditory neurons (Figure 4G). The absence of positive responses to gaps suggests the omission of a stimulus in a predictable sequence is not represented in the brain of larval zebrafish. While we did not find any relationship between the magnitude of these dips and the number of preceding stimuli (Figure S2G), it is possible that these events are encoded by suppression of neurons, and the overall elevated calcium levels in our experiment did not permit us to reliably capture suppression^49^.

Based on these results, future studies could be refined to improve the possibility of eliciting deviance-specific responses in larval zebrafish. The selection of pure tones could be improved by using loudness curves to better match the salience of sounds, as well as introducing more stimuli per block. An initial block of alternating stimuli could also be used to equally adapt both experimental frequencies before the first experimental block. Reversing the order of blocks 1 and 2 between animals could also be a useful approach, although it may introduce inter-individual differences in order effects.

### Limitations of our approach, and future prospects for addressing them

An inherent limitation of calcium imaging is the speed of the probes^50^. While in this manuscript we utilize the fastest GCaMP6 variant, calcium accumulation in cells limits our ability to detect very small or fast changes, that can be detected with techniques such as electroencephalography. It is possible that smaller interstimulus intervals would reveal stronger oddball responses, but the slow rise and decay of our probe did not permit us to investigate this possibility. This limitation is especially evident in the gap experiments, where the large drop in calcium potentially masks subtle differences in activity during this time. Regardless of the speed of the probe, calcium accumulation in cells required us to use highly stringent criteria to exclude possibly confounding results. Voltage indicators or more recent generations of calcium indicator are a promising future avenue to investigate similar paradigms with greater subtlety and faster imaging rates.

The 6 dpf larvae tested in this preparation permit whole-brain imaging at a single cell resolution, however many higher cognitive functions such as associative learning and social interaction occur at later developmental stages^51–55^. While it is difficult to match developmental stages between mammals and fishes, the MMN response in humans is not present in early infancy, and begins to resemble the adult response at around 6 months^56,57^. It is therefore possible that the SSA of responses detected here in larval zebrafish only become behaviourally relevant later in the life of the animal, and that the associated functional circuits are not yet fully established. Similarly to the mammalian cortex, the telencephalon is associated with cognition in adult fish^58–60^, but during the larval period these same regions are undergoing extensive growth, remodelling, and synaptogenesis^61^, and more complex circuitry may not yet be established. True deviance-detection responses may therefore become present in the telencephalon at later developmental stages.

Although our microscope’s field of view would not capture the entire brain of zebrafish at older developmental stages, targeted studies of individual brain regions at single-cell resolution are technically feasible^62–64^.

## Conclusions

Altogether, we report the first evidence for auditory stimulus specific adaptation in teleost fishes. While these responses are a requirement for deviance detection, we did not find definitive deviance-specific responses in larval zebrafish, nor did we find responses to the omission of a stimulus in a predictable sequence. Further studies could expand on the experiments presented here to identify the ethological relevance of frequency selectivity in zebrafish, and investigate whether true deviance-detection responses develop with age.

## Author Contributions

Experimental design: MW, REP, JBM and EKS. Experimental methodology: MW, REP, IAF-B. Data collection: MW. Processing of calcium imaging data: JA and WQ. Data analysis: MW and SJS. Writing: MW, SJS and EKS. Supervision: SJS and EKS. All authors read and approved the final manuscript.

## Acknowledgements

The authors would like to acknowledge the University of Queensland’s Biological Resources aquatics team for maintenance of zebrafish lines, and the Queensland Brain Institute workshop staff for 3D-printing of imaging chambers.

## Ethics approval

All work was performed according to research application SBS/341/19 and breeding application IMB/271/19/BREED, which was approved by the Anatomical Biosciences Animal Ethics Committee at the University of Queensland.

