## Supplemental figures for "Evidence for auditory stimulus-specific adaptation but not deviance detection in larval zebrafish brains"

SUPPLEMENTAL FIGURES & LEGENDS:

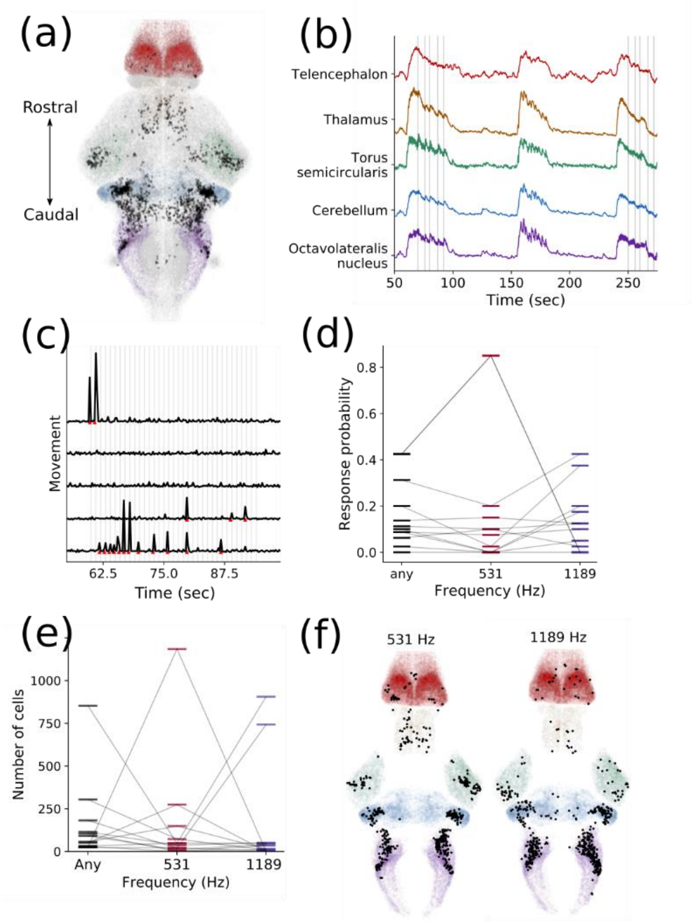


**Figure S1: Auditory supplemental information. A)** Distribution of auditory neurons, with auditory regions highlighted. The auditory neurons (black) almost entirely fall within the octavolateralis nucleus (purple), cerebellum (blue), torus semicircularis (green), thalamus (orange), and telencephalon (red). **B)** Average ΔF/F across brain regions implicated in the auditory processing pathway. Vertical lines indicate oddball tone presentations. **C)** Inferred fish movement for 5 example animals. Vertical lines represent stimulus presentations, red arrows indicate detected movement. **D)** Behavioural response probability (as inferred from motion correction) to any tone, 531 Hz, or 1189 Hz across all animals. Each line represents one fish. **E)** Contributions of each fish to generally auditory responsive cell populations (black), 531 Hz - preferring neurons (pink), and 1189 Hz – preferring neurons (purple). Each line represents a fish. All fish contribute to each group, but to different extents. **F)** Distributions of frequency-specific auditory neuron populations overlaid with outlines of auditory brain regions: octavolateralis nucleus (purple), cerebellum (blue), torus semicircularis (green), thalamus (orange), and telencephalon (red).

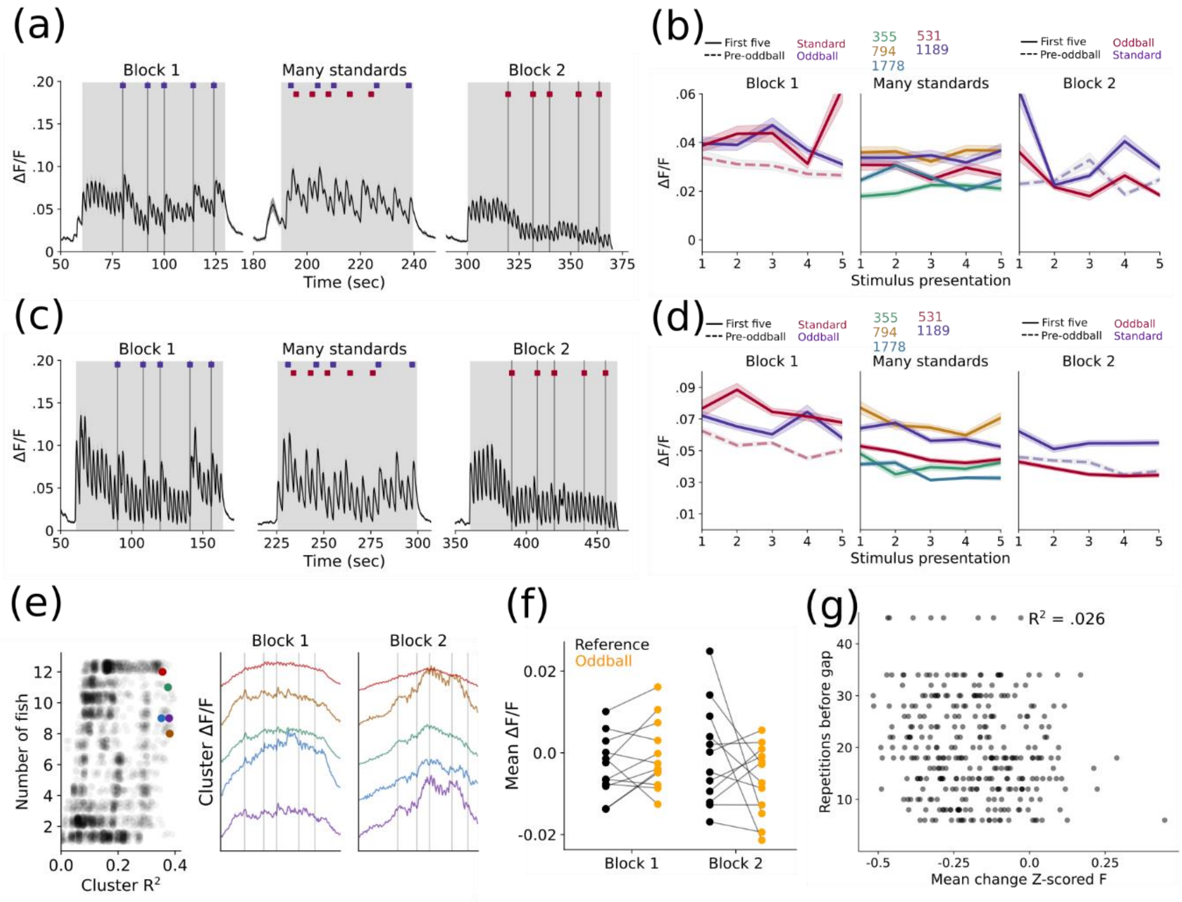


**Figure S2: Stimulus specific adaptation and oddball supplemental information. A)** Mean activity trace of all auditory responsive neurons in a dataset with a 2 second interstimulus interval. **B)** Mean response amplitudes of all auditory-classified neurons to specific stimulus presentations in a 2 second interstimulus interval dataset. In block 1 and 2, solid lines represent the first 5 presentations of each frequency (as either standard or oddball), and the dotted line represents the 5 presentations of the standard stimulus directly preceding an oddball. In the many standards block, the mean responses to 5 presentations of each frequency are shown. Shading represents standard error of mean. **C-D)** Same as A and B but for a dataset with a 3 second interstimulus interval. **E)** Left: clusters identified by 100 permutations of k-means clustering with 30 clusters, according to their representation across fish and correlation to the oddball stimulus. Right: Mean traces of example clusters highlighted on the right with high correlation to oddball stimuli. These activity traces do not resemble oddball-specific responses. **F)** Comparison of amplitude of responses from neurons in pooled clusters with $R^2 > 99^{th}$ percentile to oddball stimuli (yellow) and preceding standard stimuli (black) in each block. **G)** Mean ΔF/F by number of preceding sounds in the silent gap dataset. The magnitude of the dip at the time of the gap is not affected by the number of preceding sounds ($R^2$ = 0.026).
